# Perceived and observed biases within scientific communities: a case study in movement ecology

**DOI:** 10.1101/2024.07.29.605602

**Authors:** Allison K. Shaw, Leila Fouda, Stefano Mezzini, Dongmin Kim, Nilanjan Chatterjee, David Wolfson, Briana Abrahms, Nina Attias, Christine E. Beardsworth, Roxanne Beltran, Sandra A. Binning, Kayla M. Blincow, Ying-Chi Chan, Emanuel A. Fronhofer, Arne Hegemann, Edward R. Hurme, Fabiola Iannarilli, Julie B. Kellner, Karen D. McCoy, Kasim Rafiq, Marjo Saastamoinen, Ana M. M. Sequeira, Mitchell W. Serota, Petra Sumasgutner, Yun Tao, Martha Torstenson, Scott W. Yanco, Kristina B. Beck, Michael G. Bertram, Larissa T. Beumer, Maja Bradarić, Jeanne Clermont, Diego Ellis-Soto, Monika Faltusová, John Fieberg, Richard J. Hall, Andrea Kölzsch, Sandra Lai, Larisa Lee-Cruz, Matthias-Claudio Loretto, Alexandra Loveridge, Marcus Michelangeli, Thomas Mueller, Louise Riotte-Lambert, Nir Sapir, Martina Scacco, Claire S. Teitelbaum, Francesca Cagnacci

## Abstract

Who conducts biological research, where, and how the results are disseminated varies among geographies and identities. Identifying and documenting these forms of bias by research communities is a critical first step towards addressing them. We documented perceived and observed biases in movement ecology. Movement ecology is a rapidly expanding sub-discipline of biology, which is strongly underpinned by fieldwork and technology use. First, we surveyed attendees of an international conference, and discussed the results at the conference (comparing uninformed *vs* informed perceived bias). Although most researchers identified as bias-aware, only a subset of biases were discussed in conversation. Next, by considering author affiliations from publications in the journal *Movement Ecology,* we found among-country discrepancies between the country of the authors’ affiliation and study site location related to national economics. At the within-country scale, we found that race-gender identities of postgraduate biology researchers in the USA differed from national demographics. We discuss the role of potential specific causes for the emergence of bias in the sub-discipline, e.g. parachute-science or accessibility to fieldwork. Undertaking data-driven analysis of bias within research sub-disciplines can help identify specific barriers and first steps towards the inclusion of a greater diversity of participants in the scientific process.

## Introduction

Biases (systematic distortions with respect to the distribution of a reference population) universally affect the production and dissemination of scientific research. These biases include the topics prioritised for funding [1,2] and publication [3,4], which papers are cited [5–7], the language in which results are communicated [8,9], and the identities of authors in the peer-reviewed literature, especially in high-impact international journals [10,11]. Indeed, the composition of the scientific community itself reflects long-standing biases in terms of gender [12–15], race and ethnicity [16,17], socioeconomic status and family education level [18,19]. Beyond ethical concerns, these biases limit scientific progress and constrain insights and innovations [20]. For example, a lack of researcher gender diversity can limit the research questions and topics addressed [21]. Cultural biases can also impact interpretation of science [22]. Finally, omission of Indigenous Knowledge can lead to knowledge gaps: traditional sustainable ways of living [23], or lost animal migratory corridors [24], and perpetuate inequalities that impact science policy decision-making [25]. Within biology, biases also include the geographic locations of research [7,26] and taxonomic group(s) studied [27–29].

There is an urgent need for the scientific community to document how these biases develop, persist, and change, in order to work proactively towards the goal of broadening equitable participation [30]. Yet, the range of these biases are rarely quantified within specific disciplines. Only studying biases at a broad disciplinary level (e.g., across all of biology) can risk overlooking important sub-discipline-specific factors which are needed to formulate targeted actions to rectify inequities. Aiming to solve issues at the sub-discipline level can be more approachable. Often in biology, there are sub-discipline-specific communities that can be succinctly targeted to improve their participation on shorter time scales than an entire disciplinary field. Case studies within sub-disciplines can also illustrate concrete examples of biases and suggest potential paths forward. Within a sub-discipline, examining bias at different scales (for example, clarifying if biases manifest within or among countries; [9]) helps shape the scope and nature of potential solutions. Furthermore, determining the extent to which researchers in a sub-discipline discern biases (perceived bias) and the degree to which biases manifest in the quantifiable activities of that discipline (observed bias) represent critical initial steps in identifying potential solutions.

This paper uses movement ecology as a case study of bias across scales. Movement ecology [29,31,32] is an emerging sub-discipline of biology that is increasingly represented in high-impact journals [33–35], and is the subject of a recently launched journal (*Movement Ecology*), and multiple international conferences. The recent growth of this sub-discipline emphasizes the need to critically examine and address its embedded biases, as it continues to grow. The fieldwork-intensive nature of movement ecology often requires expensive technology, large datasets, remote travel, specialised training, and computational skills and resources, all of which can magnify extant biases.

Here, we (a biased sample of the movement ecological community; see supplement 1) describe biases present within movement ecology. Our overall goals are to start a conversation on rectifying bias in our community, and to present a case study that can be used by other sub-disciplines within biology. We use four approaches to consider both perceived biases (looking at uninformed and informed perspectives) and observed bias (considering two forms of bias at two spatial scales). First, we quantify perceived biases by surveying attendees of a conference in our sub-discipline. Next, to quantify the extent to which biases manifest, we assess data on observed biases at two scales: among countries and within a specific country. Our among-country observed bias approach quantifies patterns in countries where authors are based and where research is conducted for articles published in the sub-discipline’s primary journal. Our within-country observed bias approach quantifies representation in the broader discipline of biology with respect to race and gender identities of academic biologists in the United States. We then summarise a conference-based discussion explicitly organised around these three findings (survey results, international authorship patterns, within-country identity patterns), i.e., the perceived bias informed by observed bias data. Finally, we contextualise the results within our sub-discipline, identifying traits potentially driving the emergence of specific biases.

## Methods

### Uninformed perceived bias: Pre-conference Survey

To quantify perceived bias within the movement ecology community, we conducted an anonymous email survey of attendees registered for an international conference in the sub-discipline. The conference, held in Tuscany Italy in May 2023, was the third edition of a thematic conference series on movement ecology of animals that primarily addresses an audience of specialists in the sub-discipline. The attendees of these conferences are typically a mixture of invited speakers and selected poster presenters, with diversity being a criterion of selection alongside scientific excellence as per the conference series guidelines. The conference participation is capped at 200 individuals; in 2023 it was attended by 196 people from 24 countries (2 countries in Africa, 5 in Asia, 2 in Oceania, 13 in Europe, 2 in North America; for demographics over time see figure S1 in supplement 2). The conference fee exceeded US$1300 (including accommodation and food, but not transportation). Some of this cost was offset for some participants by conference grant funding. The survey was distributed via Qualtrics, an online survey platform that meets human subjects research data security requirements (see supplement 3). We used a mixed approach with both short-answer questions (harder to analyse quantitatively but open-ended responses) and multiple choice questions (easier to analyse quantitatively, but constrained responses). We asked registered attendees of the conference to what extent they perceived bias in the community (Q1), what the main sources of bias were (Q2 short answer, Q3 multiple choice), which experiences their answers were based on (Q4), how the community could become less biased (Q5), and who should be responsible for addressing bias in the community (Q6). To generate the possible answers for Q3 on probable sources of bias, we requested input from researchers of different backgrounds who work on movement ecology but were not attending the conference. The study protocol was reviewed by the University of Minnesota Institutional Review Board (IRB, which assesses the safety of research with human subjects) in March 2023 and was determined to be exempt from further IRB review. We distributed the survey to approximately 200 registered attendees via email on April 25th 2023 (with a reminder on May 10th 2023). A total of 135 survey responses were collected between April 26th and May 24th 2023, prior to the conference. Survey responses were analysed quantitatively for Q1, Q3, Q4 and Q6, and qualitatively for the open-ended questions (Q2, Q5).

### Observed bias

#### International scale, geographic bias: Journal articles

To provide an example of observed bias (in researcher representation) at the international scale, we analysed geographic patterns of authors’ institutional affiliation and study location for articles published in the open-access, flagship journal of our sub-discipline, *Movement Ecology.* On January 21st 2023, we used Scopus to download article information (including the affiliations of all authors and the address for correspondence) for all papers published in this journal to date. There were 370 articles published between the journal’s inception in 2013 to the data download date. For each article, we extracted (i) the first country listed for the first author (i.e., their first affiliation if more than one was listed), and (ii) the first country listed for the corresponding author. We quantified the number of times each country appeared in each of these lists (first author or corresponding author) for the full set of 370 articles, and looked at the distribution among countries. For each article, we also determined whether the primary data used in the article was newly collected for the study and, if yes, the country of data collection. For studies that tracked animals that crossed international borders, we only included the country/territory where the trackers were deployed. Of the 370 articles, 266 were considered to have collected new data, while the other 104 were excluded from this part of the analysis because they were corrections, review papers, meta-analyses, theoretical, or that used simulated or previously published data. For consistency, we used country/territory names as they appear on the World Bank list [36]. We used World Bank data [37] to get the most recently available yearly Gross Domestic Product (GDP) data for each country where primary authors were based. We then estimated the effects of yearly GDP on each country’s yearly number of publications using a Generalized Additive Model [38]. The model used a Poisson family of distributions and a log link. The model included a smooth effect of log_10_(GDP) (as the distribution had a long right tail) and a random intercept of country to account for country-level over-representation (model G sensu Pedersen et al. [39]):

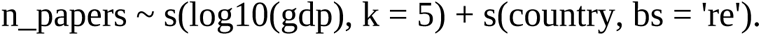

It was not possible to include random smooths for each country (model GS sensu Pedersen et al. [39]) because a third of the countries in the dataset only had publications for one year. We also used 2022 gross national income (GNI) levels from the World Bank Atlas [40] to classify countries into different income levels: low income countries have a GNI of <$1,135, lower middle-income have a GNI between $1,136 and $4,465, upper middle-income have a GNI between $4,466 and $13,845, and high income countries have a GNI over $13,846. We grouped countries into regions based on the World Bank list [36] and then created an alluvial plot, linking each study from the institutional affiliation to the location of fieldwork using the *ggalluvial* R package [41] (see also supplement 4).

#### Within-country scale: race and gender bias in USA biology researchers

To provide an example of observed bias (in demographic representation) at the within-country scale, we analysed patterns of representation by race/ethnicity and gender across career stages of academia and compared them to the general public for the United States of America (USA). We chose the USA because it is a country that collects and publicly disseminates this demographic data on a national scale. The USA is also the country with the most first authors of *Movement Ecology* articles (n=120 of 370) analysed above. Since data is not available down to the sub-discipline scale of movement ecology, we used the closest scale available: biology. We gathered 2021 data for graduate students and postdoctoral researchers [42] as well as faculty [43], and census data for the USA general population for 2019 (the most recent data available) [44]). We combined some race/ethnicity categories to facilitate comparison (see Table S2 in supplement 4). All datasets categorised gender as a binary (male or female). For each group, we calculated the proportion of individuals who identified in each gender-and-race combination (hereafter ‘race-gender group’). For the graduate student and postdoc data, race/ethnicity was only collected for individuals that are either USA citizens or permanent residents; so for these datasets we calculated the proportion of individuals in each race-gender group as a portion only of the number of USA citizens / permanent residents. For each race-gender group for each career stage, we calculated the relative representation [45] as

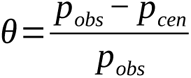

where *p*_obs_ is the observed proportion of a race-gender group within a career stage and *p*_cen_ is the proportion of a race-gender group across the censused USA population.

### Perceived bias informed by observed bias data: Conference Discussion

Finally, to assess observation-informed perceived bias, results of both the survey and the observed bias approaches were presented and discussed at the previously mentioned conference in a session on inclusion and barriers to inclusivity. The day before the session, all conference attendees were provided a handout summarising the findings of the survey and the international and within-country approaches (supplement 5) and were invited to attend the discussion. Of the 196 conference attendees, 105 participated in the discussion. Attendees were split into 10 groups of 8-12 people. The discussion volunteer leaders described the goals of the session, presented an overview of the findings, and provided groups with two sets of discussion prompts. Each group identified a scribe to record notes. The groups discussed a first set of prompts (“In your words, what do you think bias is? Before now, have you ever thought about bias in the movement ecology community? What is surprising or interesting to you about the survey results?”), and then reported back to the broader group. The groups then discussed a second set of prompts (“What is something you plan to do going forward? What should we discuss doing as a community?”) and again reported back to the broader group. Notes from each group were compiled by the discussion leaders, and summarised below, and everyone was invited to be a potential co-author on this publication (see supplement 1).

## Results

### Uninformed perceived bias: Pre-conference Survey

#### Frequency and sources of bias

We received survey responses from 135 individuals out of approximately 200 distributed surveys (although not all respondents answered every question). Nearly all survey respondents who answered question Q1 (see supplement 3), believed that there was bias present within the movement ecology community (96.8%, n = 90; Figure 1A). The majority of respondents rated this bias as low (44.1%, n = 41, scores 1-2), whereas 20.4% (n = 19) of respondents perceived bias as high (scores of 4-5). These patterns largely correlated with the respondent’s own experience of bias as identified in survey question Q3. Specifically, 55.6% (n = 25) of individuals with no direct experience of bias perceived little to no bias within the community, compared to 39.6% (n = 19) of individuals with personal experience of bias. Conversely, only 6.7% (n = 3) of individuals with no direct experience of bias perceived the community as highly biased, compared to 33.3% (n = 16) of individuals with personal experience.

**Figure 1.**
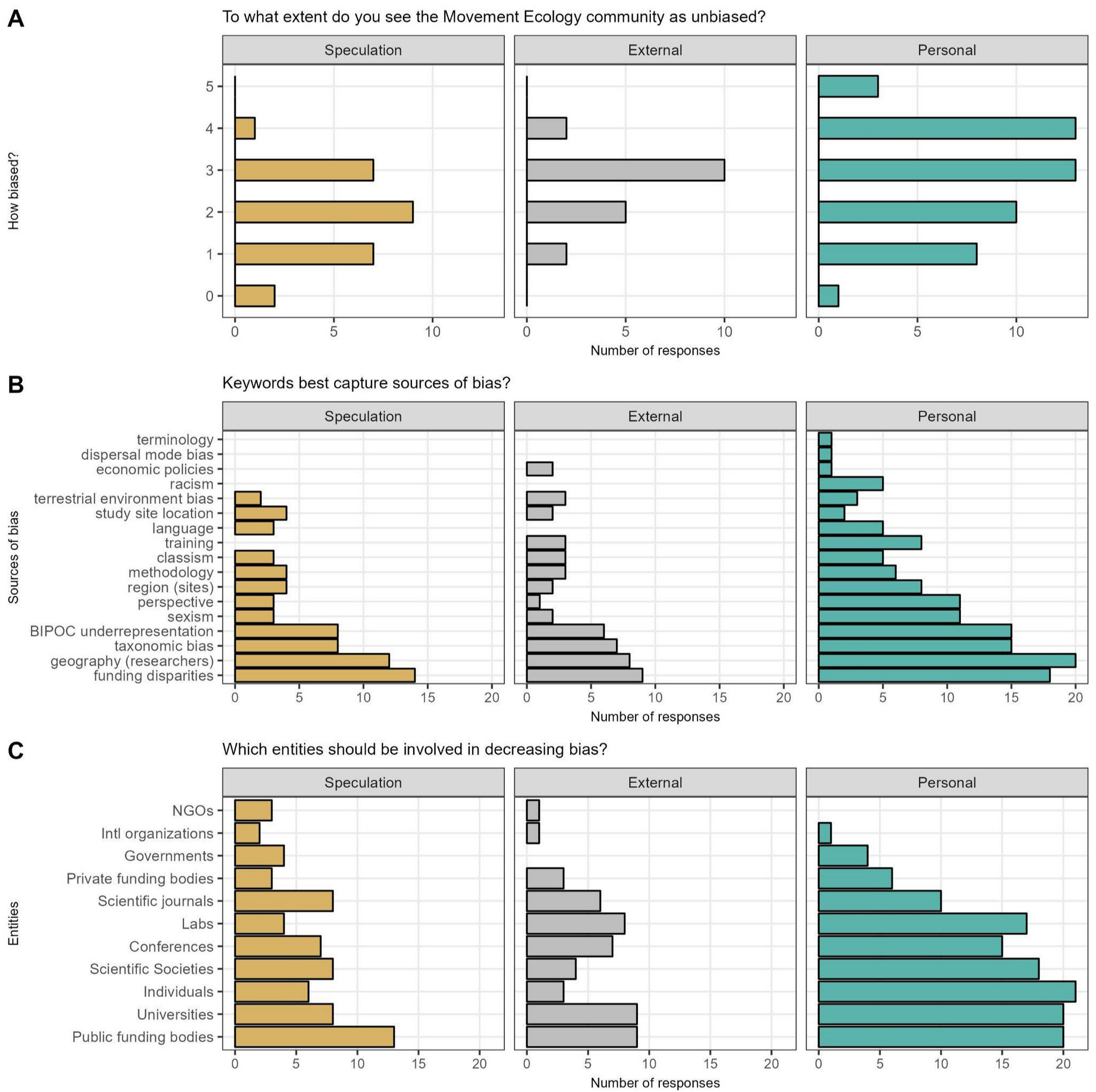
Summary of survey responses from participants of an international conference in movement ecology split by individuals’ own experiences of biases. Responses cover the questions of (A) bias within the movement ecology community (0 is unbiased, 5 is very biased), (B) sources of bias, and (C) entities that should be involved in decreasing bias. Individual’s responses were based on one of three categories: speculation (yellow), external evidence such as readings (grey), and personal experience (green).

Survey respondents’ own experience of bias influenced not only their perceived frequency of bias, but also the sources of bias. In response to the open-ended question Q2 ‘What do you think the main sources of bias are, if any?’, the most commonly reported answers were biases associated with who does the study (Table S1 in supplement 2). In response to a close-ended question Q4, individuals with personal experience of bias reported the greatest number of perceived sources of bias (n = 17) relative to those with no direct experience (n = 13) (Figure 1B). Overall, ‘funding disparities’, ‘geographic concentration of researchers’, ‘taxonomic bias’, and ‘BIPOC [Black, Indigenous, and other People of Color] underrepresentation’ were consistently reported as key sources of bias, accounting for 55.2% (n = 88) of the keywords best capturing the sources of bias (see supplement 3 for full list of options in the survey). In contrast, there were differences in perceived sources across groups with different experiences of bias. For example, explicit categories of discrimination - ‘sexism’, ‘racism’, and ‘classism’ - accounted for 15.6% (n = 21) of keywords for individuals with direct experience of bias, ranking 5th, 7th, and 8th, respectively. However, ‘sexism’ and ‘classism’ were less perceived as sources of bias by respondents with no personal experience of bias, with ‘racism’ not perceived as one of the top three issues. Together, these data highlight the prominent role of personal experience in dictating the perception of frequency and source of bias within a research community.

#### Reducing bias

Suggested strategies for reducing bias (Q5) ranged from interventions and actions targeted at the individual level, for example elevating researchers from under-represented backgrounds, to the institutional level, such as changing funding priorities (Table 1). Overall, public funding bodies, universities, scientific societies, and individuals were most frequently mentioned as the entities that should be involved in reducing bias (55.8% of keywords, n = 88, Q6) (Figure 1C). However, whilst there was consensus among groups with different experiences of bias that public funding bodies, scientific societies, and universities should be involved in reducing bias, most individuals who had not personally experienced bias did not believe that individuals should be charged with reducing bias.

**Table 1.**
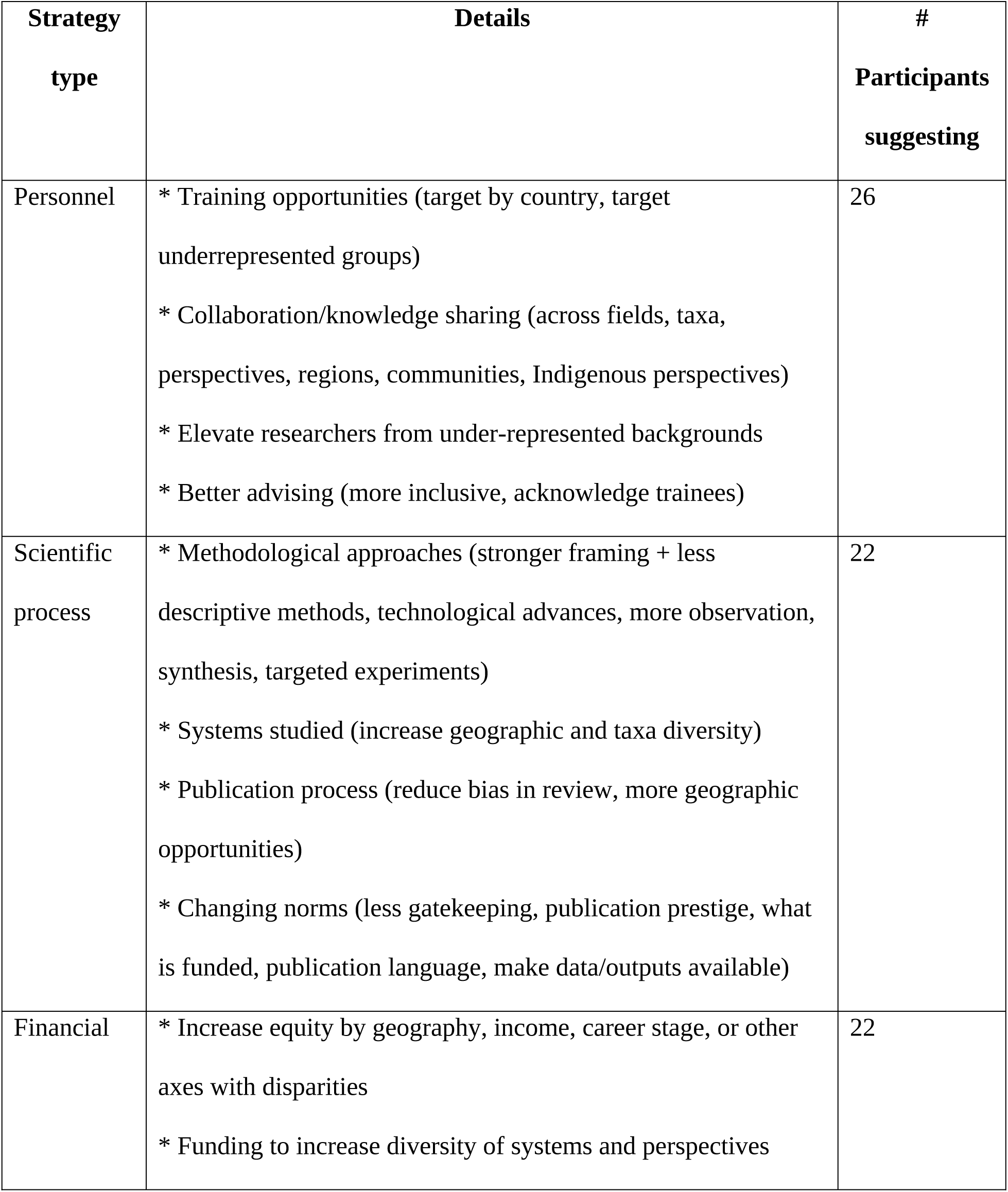
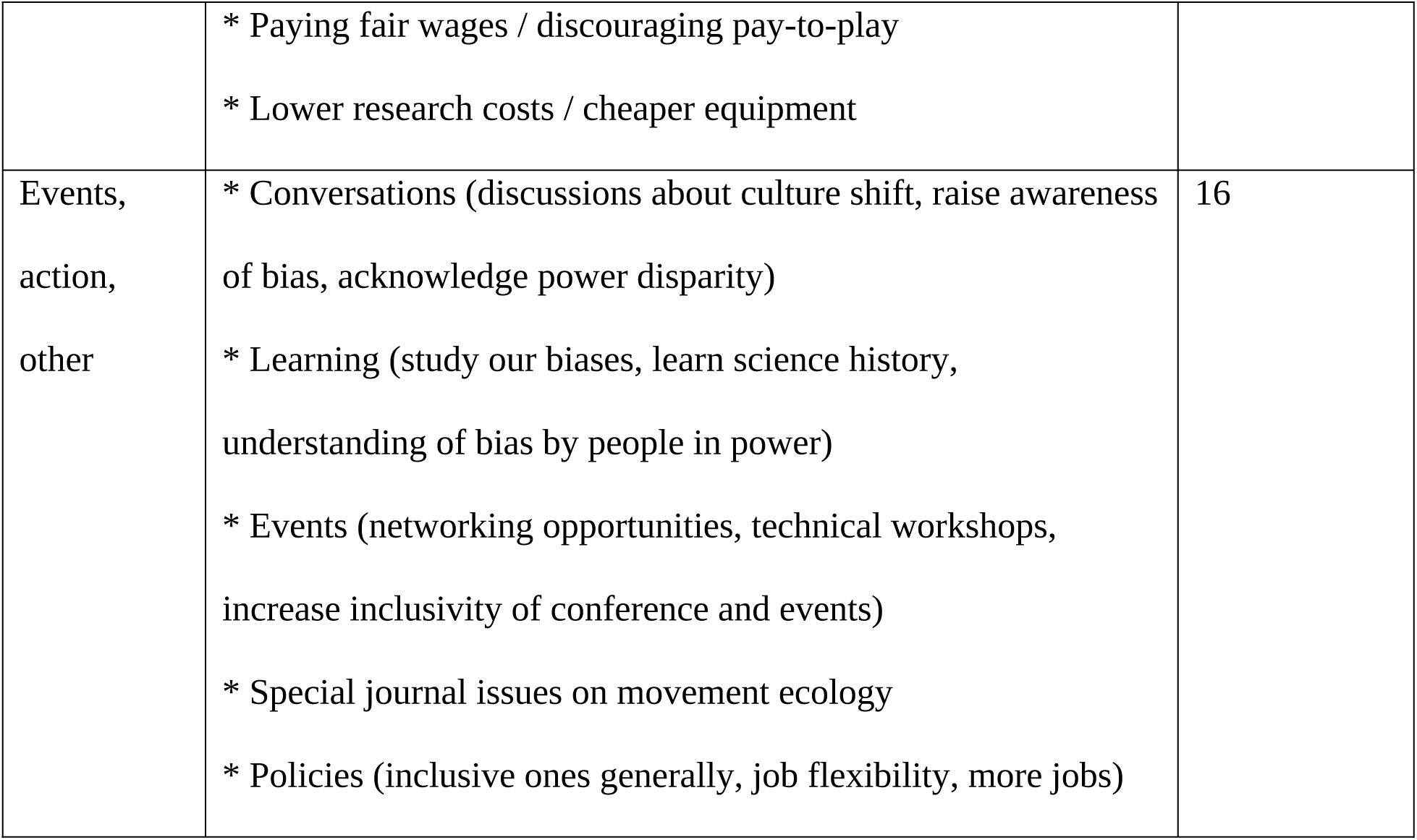
Responses to the open-ended survey question [Q5] ‘How do you think the Movement Ecology community could become less biased?’, organised into broad categories. Of 135 survey takers, 61 answered this question.

| Strategy<br>type | Details | #<br>Participants<br>suggesting |
| --- | --- | --- |
| Personnel | * Training opportunities (target by country, target underrepresented groups)<br>* Collaboration/knowledge sharing (across fields, taxa, perspectives, regions, communities, Indigenous perspectives)<br>* Elevate researchers from under-represented backgrounds<br>* Better advising (more inclusive, acknowledge trainees) | 26 |
| Scientific<br>process | * Methodological approaches (stronger framing + less descriptive methods, technological advances, more observation, synthesis, targeted experiments)<br>* Systems studied (increase geographic and taxa diversity)<br>* Publication process (reduce bias in review, more geographic opportunities)<br>* Changing norms (less gatekeeping, publication prestige, what is funded, publication language, make data/outputs available) | 22 |
| Financial | * Increase equity by geography, income, career stage, or other axes with disparities<br>* Funding to increase diversity of systems and perspectives | 22 |
|  | * Paying fair wages / discouraging pay-to-play<br>* Lower research costs / cheaper equipment |  |
| Events,<br>action,<br>other | * Conversations (discussions about culture shift, raise awareness of bias, acknowledge power disparity)<br>* Learning (study our biases, learn science history, understanding of bias by people in power)<br>* Events (networking opportunities, technical workshops, increase inclusivity of conference and events)<br>* Special journal issues on movement ecology<br>* Policies (inclusive ones generally, job flexibility, more jobs) | 16 |

### Observed bias

#### International scale, geographic bias: Journal articles

The analysis of geographic bias at the international scale showed that countries were not evenly represented in authors’ national affiliations for articles published in the journal *Movement Ecology*. Only 28 countries (<15% of global countries) were represented by first authors in the journal (n = 370 articles; Figure 2). There were 120 first authors affiliated with the United States of America, 40 with Canada, 40 with Germany, and 39 with the United Kingdom. All other countries were represented by less than 20 first authors. The patterns were similar when looking at the country of affiliation of the corresponding author (see Figure S2 in supplement 2). Additionally, the average number of publications per year increased exponentially with yearly Gross Domestic Product (GDP; an indicator of the size of a country’s economy; see Figure 3, Figures S3-S4 in supplement 2).

**Figure 2.**
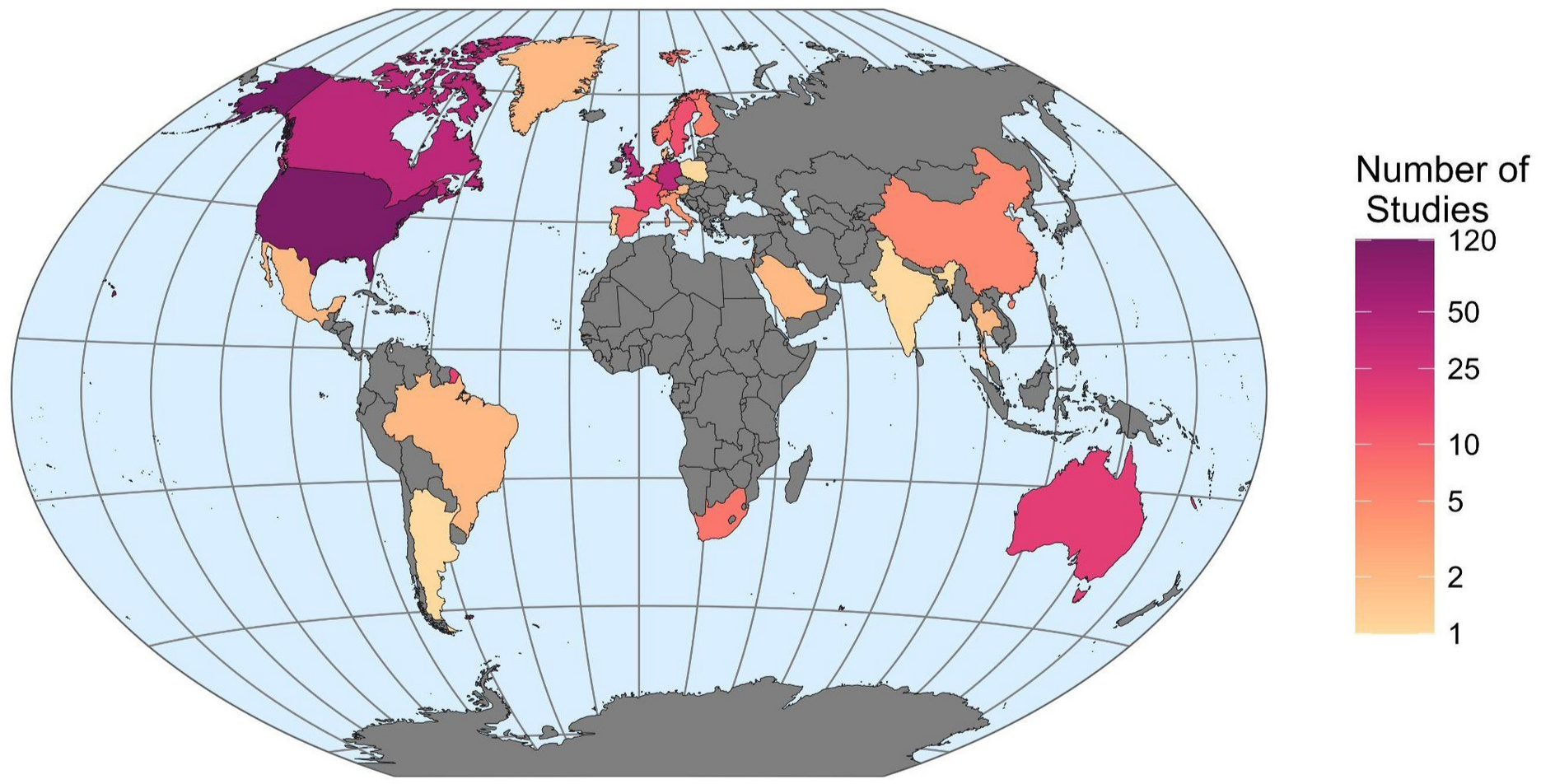
The number of times each country was listed as the first author’s first affiliation for all 370 articles published in the journal *Movement Ecology* from its launching in 2013 until January 2023.

**Figure 3.**
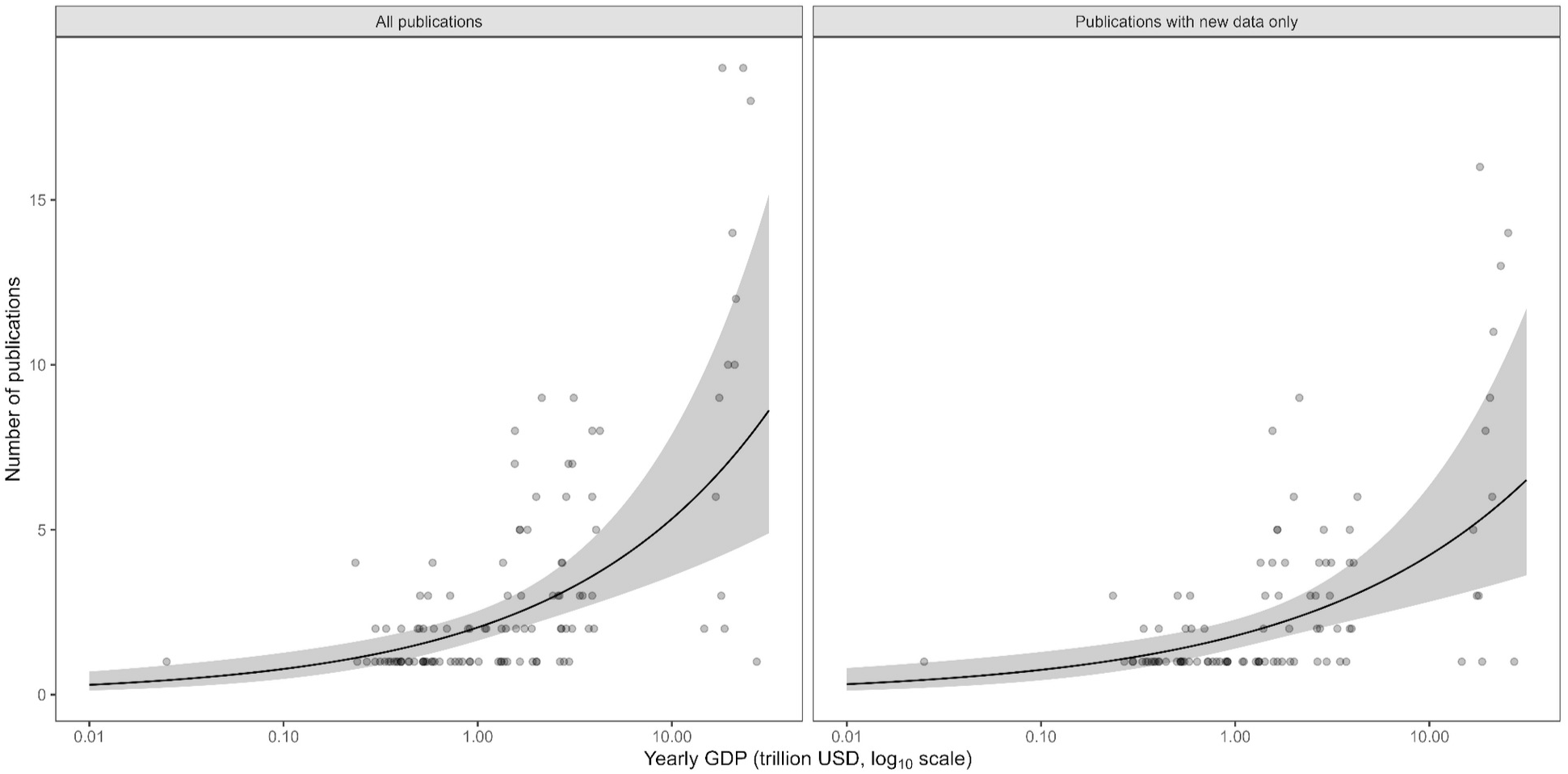
Publication rate depends strongly on a country’s gross domestic product. The relationship between a country’s GDP and the number of times it was listed as the first author’s first affiliation across (left) all 370 articles and (right) the 266 articles with new and empirical primary data, published in the journal *Movement Ecology* from its launching in 2013 until January 2023. The black line indicates the estimated relationship, while the grey shaded areas indicate the 95% Bayesian credible intervals.

Most studies were conducted in the same geographic region as the first author’s institutional affiliation. Of the 266 articles that collected new empirical data, 78% of studies conducted fieldwork in the same geographical region as the institution of the first author (Figure 4). For the 22% of the articles where the study site and first author affiliation were not identical, there were some notable disparities. First authors with institutional affiliations in North America and Europe had a higher tendency to conduct fieldwork in other regions when compared to first authors with institutional affiliations in Asia, Africa, and Latin America. Most studies in Latin America were conducted by researchers based at institutions in North America. Most studies in Africa were conducted by researchers based at institutions in either North American or Europe with a fairly even split between the two. Furthermore, most studies that took place in low and lower-middle income countries were led by researchers with institutional affiliations in other regions (Figure 4, red and green bars). For studies in upper-middle income countries, the pattern varied by region – studies in Europe and Asia were conducted by researchers at institutions in those same regions whereas studies in Africa and Latin America were conducted by researchers at institutions in other regions (Figure 4, yellow bars).

**Figure 4.**
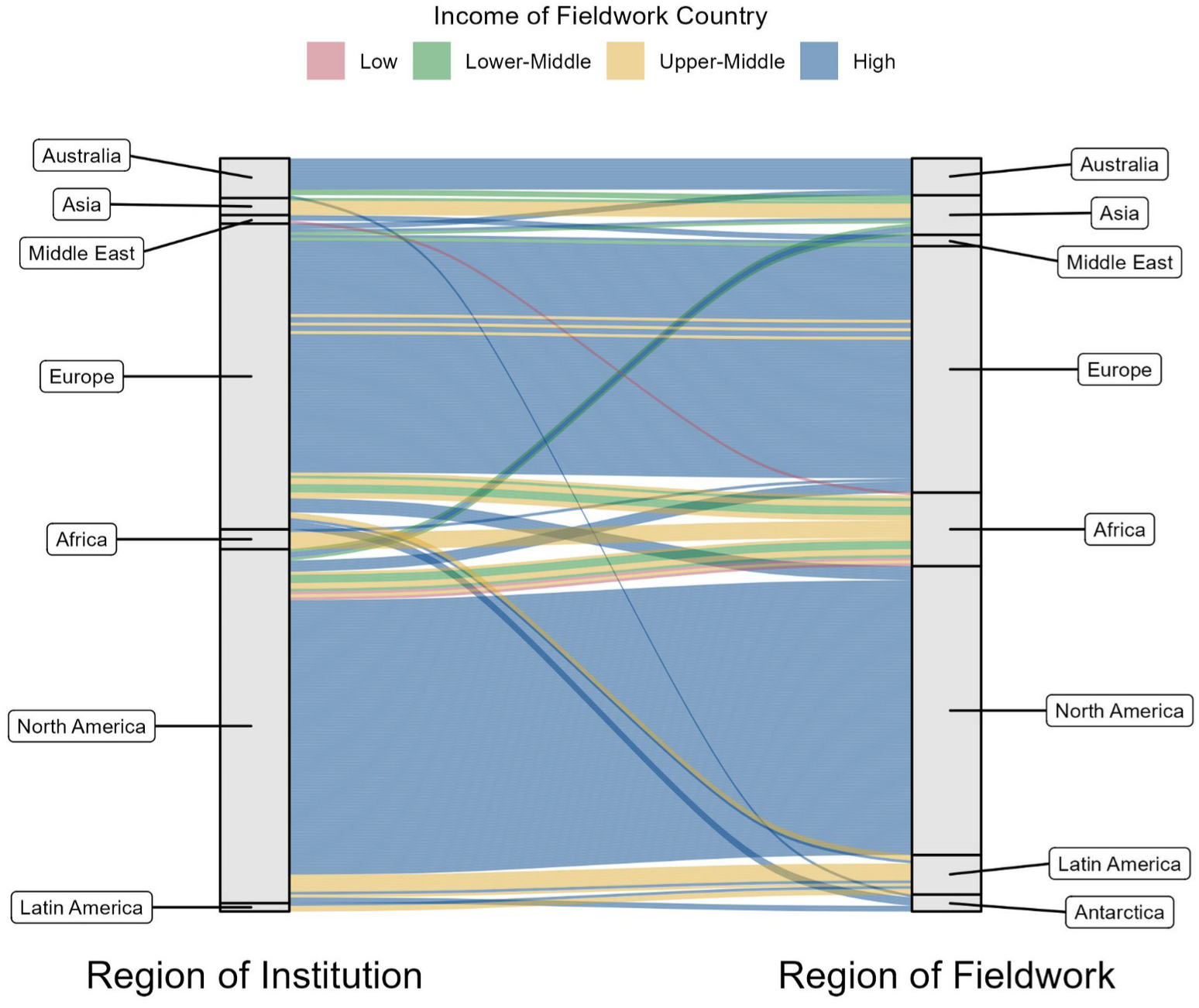
Comparison of first author’s regional affiliation (left) to region of fieldwork (right) for the 266 *Movement Ecology* studies (from 2013 until January 2023) with new empirical data. We used only the site of deployment for studies that tracked animals across multiple countries. Gross national income in 2022 (GNI) and regional categorization included in this figure are reported by the World Bank: low (GNI <$1,135), lower middle ($1,136 to $4,465), upper middle ($4,466 to $13,845), and high (GNI > $13,846).

#### Within-country scale: race and gender bias in USA biology researchers

Changing scales, we next analysed observed representational bias occurring at the within-country scale for academics within the discipline of biology in the United States of America. We found that representation of race-gender groups often differed from their representation in the general USA population (Figure 5). White-male, Asian-male and Asian-female groups were all overrepresented among biology faculty compared to the general population, while all other race-gender groups were underrepresented. The most underrepresented race-gender group at all career stages was Black / African-American men, with underrepresentation generally increasing across career stages, and always lower than Black / African-American women. Within the race/ethnicity identities of White, Asian, and Hispanic / Latino, women had higher representation than men among graduate students but lower representation at the faculty level.

**Figure 5.**
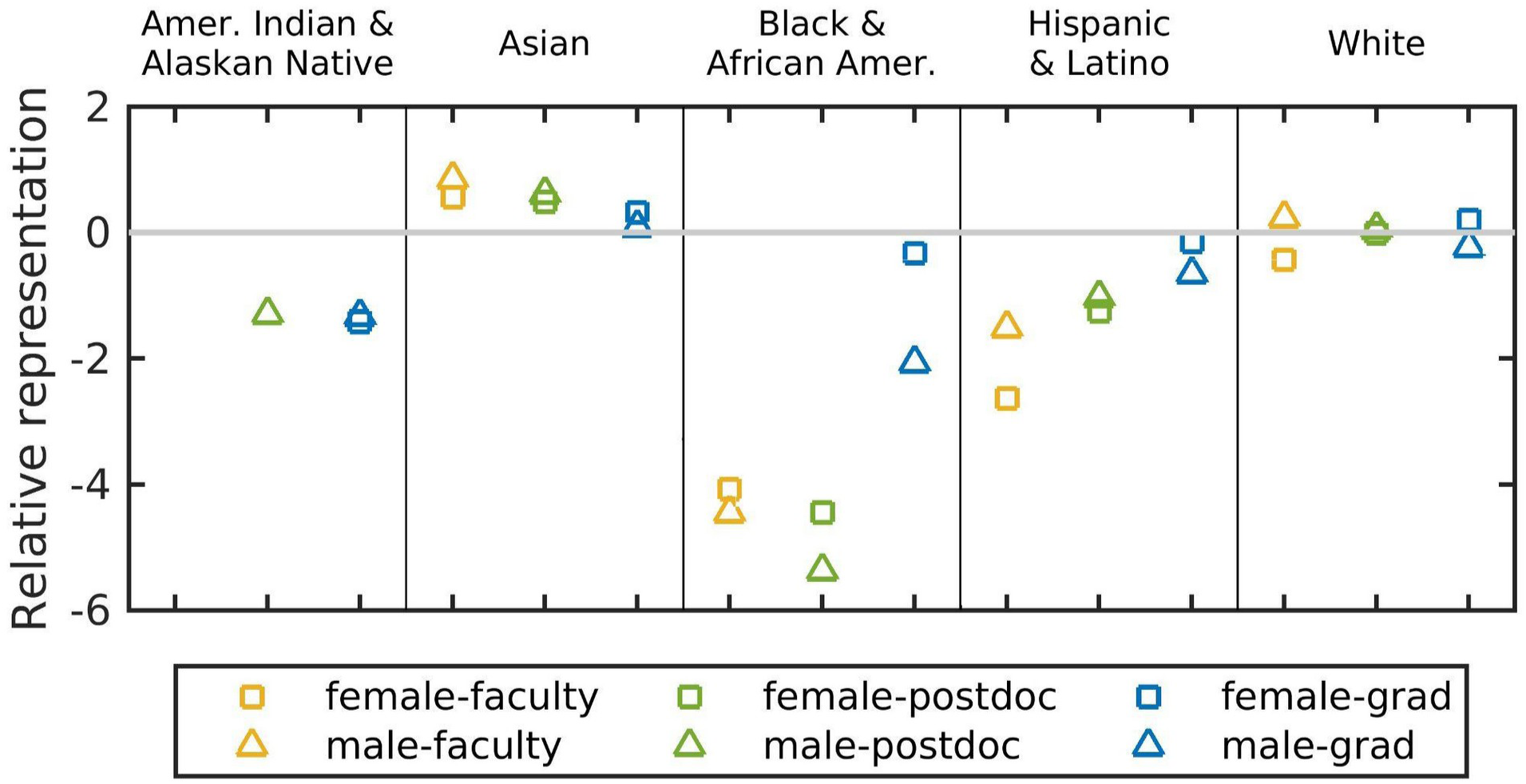
The relative representation of USA life science researchers across 3 career stages that provided race and gender information. Positive values indicate a group is overrepresented compared to the USA population (as reported in the USA census) and negative values indicate a group is underpresented. The race/ethnicity, gender, and career-stage categories included in this figure were specified in the surveys and census. Race/ethnicity categories included American Indian and Alaska Native, Asian (including Pacific Islander), Black and African American, Hispanic and Latino, and White. Note however that data on American Indian and Alaska Native faculty were not reported. Gender categories included only female (squares) and male (circles). Data available on career-stages included faculty (yellow), postdoctoral researchers (green) and graduate students (blue, but were only reported for USA citizens and permanent residents).

### Perceived bias informed by observed bias data: Conference Discussion

#### In your words, what do you think bias is?

When asked to define bias in their own words, the participants gave a variety of answers that described bias as an intentional (explicit, conscious) or unintentional (implicit, unconscious) show of prejudice that can lead to systematic disparities in representation or opportunity. Bias can encompass both outcomes and causes – both aspects emerged in the discussion although not explicitly presented in the survey. Participants discussed that bias can result from misconceptions and subjective viewpoints that are not supported by evidence, and may lead to differential treatment of some individuals. People might be focused on their own ideas, perspectives and experiences, tending to associate with other people who share similar identities or similar circumstances (e.g., socioeconomic status). Further views underlined that bias can also result from power imbalances (e.g., differences in academic rank), or confirmation bias. As a result, other legitimate viewpoints arising from different experiences or values are either ignored or rejected. Biased perspectives limit opportunities for some individuals, and can therefore result in reduced representation, leading to overall reduced diversity. These disparities can have consequences that manifest in a variety of ways, from psychological well being (e.g,. feelings of unworthiness or imposter syndrome) to support for research (e.g., limited access to funding).

#### Before now, have you ever thought about bias in the movement ecology community?

Effectively all discussion participants reported having thought about bias within our community. The specific biases on which discussions focused spanned demographic, geographic, and methodological. The community is aware of demographic biases that exist among its members. The conference attendees, the majority of whom are from (or are working in) Western nations (Figure S1 in supplement 2), acknowledge that the sub-discipline continues to be dominated by white men, and that possibilities for career progression are not distributed equitably across gender and race identities. A significant point of concern was geographical bias due to the shaping of ideas, research paradigms, and methodologies being predominantly driven by scientists in North America and Europe. The costs associated with publishing in open-access journals, attending international conferences, and utilising advanced research technologies (e.g., bio-logging devices) were thought to be major contributors of geographic bias through their disproportionate impacts on researchers, research topics, and methodologies (e.g., sample size) from countries with lower GDP. An additional concern was the prevalence of “parachute or helicopter science” where researchers from these regions descend on regions with lower GDP, often sidelining local scientists, failing to give credit to local communities and missing opportunities for skill transfer, local capacity building, and knowledge exchange. Regarding research itself, biases affect methodology and topics. For example, research frequently favours popular or ‘charismatic’ species, while neglecting less well-known ones.

#### What is surprising or interesting to you about the survey results?

Multiple groups remarked on how closely the outcomes of the survey aligned with their expectations. Many of the groups discussed geographic biases underlying these results and were interested in how the data on the affiliations of first authors in the journal *Movement Ecology* would differ from the locations of data collection or the authors’ countries of origin. [Based on this discussion point, we extracted more information from the articles to make this comparison – described above, and presented in Figure 4]. Some noted that the emphasis on geographic bias might be due to the composition of the survey pool, since many respondents referenced their own personal experience as evidence.

#### What is something you plan to do going forward?

Suggestions for actionable next steps ranged from changing individual perspectives, to exerting pressure to change community culture, to making needed policy changes to the academic system. Changing individual perspectives might require taking responsibility to acknowledge and mitigate one’s personal biases, identifying power dynamics, understanding the needs and the perspectives of others, and acting on bias when witnessed. Several participants mentioned that it is hard to fight bias on their own, suggesting the need for collective action by peers and colleagues. Ideas for changes in behaviour differed across academic stage. More senior researchers indicated the need to involve people from the global community, to increase mentorship of underrepresented communities, and to increase exchange opportunities for international students to obtain experience and training and to grow their professional networks. More junior researchers suggested the need to identify and reduce geographic and gender bias in citations used in academic publications. More outreach is needed to show opportunities that are available in the sub-discipline of movement ecology, and it is important to start diversity and equity groups (including reading groups) to raise awareness of bias in our sub-discipline. For peer-reviewing, considerations on the quality of the English writing should be separated from main comments (most journals offer a section for that), and constructive comments should account for potential sources of bias authors from underrepresented groups could be exposed to. Discussions highlighted that conference attendees were mostly native English speakers. It was also underlined that cheaper, more inclusive conferences are needed to promote change (e.g., by hosting conferences in the Global South or by offering formative opportunities such as workshops). While some solutions, such as waiving conference or publishing fees, have sometimes been implemented, they are largely financial in nature and do not adequately address the full scope of the biases identified. For example, past efforts to increase the diversity of conference attendees by making funding available have fallen short (i.e. not many applications for conference scholarships from members of underrepresented groups are submitted). It was also noted that although each suggested action addresses a small part of the problem, together they will hopefully collectively improve inclusiveness within the community.

#### What should we discuss doing as a community?

When participants were asked what we should discuss doing as a community in response to the problems associated with bias in the sub-discipline of movement ecology, solutions were proposed that can broadly be categorised as relating to funding agencies, publishing, conference organisation, and community environment. That said, in many groups, there was frustration about the inefficacy of anti-bias efforts in creating desired change. The issue of funding was discussed at length, including limitations associated with funding agencies, peer-reviewed journal costs and conference fees. For example, geographic restrictions are often imposed by funding agencies, by which only people from a certain country can access funds. For instance, national funding to support conference attendance may sometimes only be used for scholars affiliated with institutions in that country. Several groups suggested encouraging senior researchers to leverage their role to do anti-bias work, such as by using relationships with tagging companies to secure more funding for scholarships. In terms of peer-reviewed publications, some comments were made about limitations associated with research costs and to what extent this should be accounted for by reviewers. Other solutions included the publication of abstracts in peer-reviewed papers in languages other than English, double blind peer-review, focusing on inclusivity in invitations to publish in journal special issues and increased accessibility of data, code and tutorials. Similarly, choosing more affordable conference venues, considering visa issues, or supporting virtual attendance may promote broader representation. Additionally, increasing the availability of funding targeted at increasing attendance by underrepresented groups could be beneficial. It was also recommended that the perception of exclusivity should be reduced, that rules around the diversity of chairs, speakers and panelists should be evaluated, and that each specific conference should rotate locations to increase accessibility from a greater diversity of geographic regions. Others suggested that in order to ensure that underrepresented communities are not disproportionately burdened with solving problems of bias, people from majority groups should increase their engagement in anti-bias work. Other proposed solutions included changes in attitudes, such as more listening and learning, i.e. making spaces more welcoming for underrepresented groups to speak up and be heard. It was also recommended that for future conferences in our sub-discipline we bring in a professional with expertise in diversity and equity to suggest actionable changes.

## Discussion

Here, we studied bias in a scientific community (i.e. science of science; [46]), mainly by characterising sources of observed and perceived bias within the sub-discipline of movement ecology as a case study. Our purpose was to explicitly bring a topic that is often implicit to the forefront, by quantifying patterns, understanding what factors shape conclusions on these aspects within our community, and providing motivation to broaden participation and inclusiveness in our sub-discipline. We quantified **observed bias** at two spatial scales and in two different forms: among-country discrepancies in where authors publishing in *Movement Ecology* are based and where data are collected, and within-country discrepancies in race/ethnicity and gender identity of biology academics compared to the general public of the USA. The first set of findings aligns with parachute science [47], which although sometimes supported by alleged good intentions, can fuel bias when not transformative beyond the single research project [48]. Also, it may indicate GDP-skewed research investments on this particular discipline. This second set of findings recovers some previously described results with respect to single axes of identity; i.e., that the representation of women decreases across academic stages and that Black / African-American and Hispanic / Latino identities are consistently under-represented within academia in the USA [12,45]. However, our findings also highlight places where considering multiple axes of identity together can provide new insights – i.e. that scale matters. Thus, care should also be taken in extrapolating our findings across scales; the within-country dataset is simultaneously more narrow (within the USA vs across countries) and more broad (all of biology vs movement ecology) than the first dataset, and biases present at one scale may not be present at others (e.g. [49]).

We examined **perceived bias** and found that it differed by context. For example, all conference discussion participants reported having previously thought about bias in movement ecology (informed perceived bias), while our pre-conference survey showed a wide range of perceived degree of bias (uninformed perceived bias, Figure 1A). Many of the themes raised in our discussions are aligned with those that have emerged in other discussions of STEM subdisciplines underpinned by fieldwork research [50,51]. Furthermore, the same sources of bias were not highlighted in both the survey and discussion. For example, both focused on geography, but the informed discussion focused less on race and gender. Specifically, for the open-ended survey question, the most common responses were gender/sexism and race/ethnicity/racism as key sources of bias (Table S1 in supplement 2). However, for the multiple choice survey question, survey takers did not select ‘racism’ as a key source of bias, but they did select ‘BIPOC underrepresentation’ and ‘geographic concentration of researchers’. Survey participants who did select ‘racism’ as a key factor were exclusively those who drew on their personal experience to answer the survey (Figure 1B). During the discussion (informed by the survey data), this pattern on sources of bias was amplified with little (if any) discussion of racism but extensive discussion of geographic and socioeconomic factors. One interpretation of these findings is differential comfort in discussing different sources of bias, especially in group settings. However, we recognize that perception shapes discussions. Thus, our discussion of perceived bias was shaped by the participants (supplement 1); a different set of movement ecologists may have had a very different discussion about bias.

Our quantitative analysis of observed and perceived bias generated a number of ideas for future quantitative research on bias. First, one could use a different classification scheme or look at a different spatial scale to consider the link between economic activity and publications. Further research should explore the disconnect we discovered between country affiliation of researchers and countries where the research is done. Although we focused on first and corresponding authors, one could test whether the patterns change by considering affiliation of senior authors. For papers with more than two authors, one could examine the diversity of middle authors’ affiliations, and whether it matches the affiliation of the first and last authors. This should be done with care as a higher diversity of countries of affiliation could be seen as positive (i.e. promoting international collaborations) or negative (i.e. relegating researchers who support local fieldwork as middle authors, rather than as leads). Some authors also had several affiliations from countries with different geographic and economic contexts, leaving unclear what portion of the research (e.g., data collection/analysis, PhD awarding, financial support) happened in which country. Second, future work could quantify biases in other aspects of identity that were not addressed by our work, especially ones where safety and physical constraints can impede travel (e.g., LGBTQIA+ identity, disability, caregiving).

The approach we used here could also be applied to other disciplines such as archaeology, geology, and anthropology that obtain data with fieldwork. We could also compare our findings with other sub-disciplines of biology to see where biases are similar and different. For example, movement ecology may have more financial driven bias than some sub-disciplines (e.g., by requiring expensive equipment, logistics and travel) or less than others (e.g., those that rely on large infrastructure such as genomics labs). Further, fieldwork can have differential impacts on researchers based on the researcher’s identity [52], the interaction with stakeholders [53], or the cultural context.

Quantifying bias does little to move the conversation forward without a call to action. A shared comment during the conference discussion regarded the need for proactive measures to elevate voices and engage groups, so that people with all perspectives/backgrounds are present where science is being discussed and conversations on how to strengthen our communities are taking place. Survey takers also suggested a number of actionable solutions (Table 1). A core part of addressing these biases is financial: the distribution of research funds is unequal (both globally and within countries) and exacerbates existing biases. Some of these are norms that can be addressed within our community (e.g. discouraging pay-to-play opportunities, paying researchers fair wages, exploring cheaper research approaches) while others require systemic change (e.g. global redistribution of funding towards underrepresented groups). Some actions could benefit from bringing in researchers from sociology, psychology and other specialised disciplines to provide structured recognition and mitigation of bias. For example, future work could consider how different forms of bias may affect one another or be correlated, and so contemporarily emerge.

## Conclusion

Bias is omnipresent in science and we need to understand how biases develop and persist in order to proactively broaden representation. One advantage of our data-driven, community-based approach is that we can now monitor the impacts of this work at the community level (i.e., if and to what extent these conversations have been improving our community since their onset). While presenting a partial view on bias in our community, we were able to identify some critical starting points that are particularly relevant to the sub-discipline (e.g., geographic and economic bias). While identification of concrete actions to address these drivers is at its infancy, we see a proactive attitude towards the risk of bias to be part of the solution.

## Funding

We acknowledge funding support from the following sources. Any opinions, findings, and conclusions or recommendations expressed in this material are those of the author(s) and do not necessarily reflect the views of the National Science Foundation, the European Union, the European Commission, or the other funders.

| Author | Source |
| --- | --- |
| AKS | US National Science Foundation grant DEB-1947406 |
| AL | COVID-19 Bio-Logging Initiative, funded in part by the Gordon and Betty Moore Foundation, GBMF9881 |
| AMM<br>S | an ARC DP DP210103091 and a Pew Fellowship in Marine Conservation by the Pew Charitable Trusts |
| CEB | LJMU School of Biological and Environmental Sciences Research Development Fund |
| DES | NASA FINESST 80NSSC22K1535 |
| EAF | the Agence Nationale de la Recherche No.: ANR-19-CE02-0015 |
| FC | NBFC to Fondazione Edmund Mach, funded by the Italian Ministry of University and Research, PNRR, Missione 4 Componente 2, “Dalla ricerca all'impresa,” Investimento 1.4, D.D. 1034 17/06/2022, Project CN00000033 |
| JBK | The Independent Research/Development Program at the National Science Foundation |
| KBB | German Research Foundation (DFG Eigene Stelle grant: 515649177) |
| KLB | National Science Foundation Virgin Islands Established Program to Stimulate Competitive Research VI-ESPCoR |
| KR | Washington Research Foundation |
| LLC | project NetCost ANR-17-CE03-0003 grant and project MUSE2018-EbOHEALTH, grant number ANR-16-IDEX-0006 |
| MGB | The Swedish Research Council Formas 2020-02293 |
| MM | The European Union's Horizon 2020 research and innovation programme under the Marie Skłodowska Curie grant agreement (101061889) and the Swedish Research Council Formas (2022-00503) |
| NC | the National Aeronautics and Space Administration award 80NSSC21K1182<br>FWF-START Y-1486 (PS) |
| SAB | Canada Research Chair Program |
| SM | UBCO Graduate Student Travel Grant |
| SY | Max Planck - Yale Center for Biodiversity Movement and Global Change |

## Acknowledgements

The co-authors would like to thank all those who took the pre-conference survey and/or participated in the in-person meeting that constitute the backbone of the empirical analysis of this manuscript.

## Data statement

The R code and data used to fit the model in figure 2 can be found on GitHub at https://github.com/StefanoMezzini/grc-2023-gdp-papers/

