## supplement 1 for "Perceived and observed biases within scientific communities: a case study in movement ecology"

### SUPPLEMENT 1: POSITIONALITY STATEMENT

This paper includes 48 co-authors; of these 44 anonymously provided some or all demographic information for this statement (some chose to skip the survey entirely or some questions within the survey). We the authors are a group of people with higher education training (n=39 authors have a PhD as their highest degree, 3 bachelors, and 2 masters). We are also primarily currently based at a University (n=30) with others at a government institution (n=5), non-academic research centre (n=4), non-profit (n=3), or research institution (n=2). We are a mix of primarily early-career (n=23) and mid-career (n=12) academics, with fewer full professors/PIs (n=6), applied scientists (n=2), and science advisors (n=1). The majority (n=39) of us have lived or worked in more than one country, with few having studied or worked in a single country (n=5). Most of us are currently based in Europe (n=25) or North America (n=16), and n=3 elsewhere (Australia, Brazil, Israel). In terms of identity, most of us have never been in a minority racial/ethnic group throughout our academic training (n=26), with many sometimes (n=14) and few always (n=3) having been in the minority. Most of us are female (n=28), and many (n=15) male. We are a mix of primarily 35-44 year-olds (n=21), and 25-34 year-olds (n=15), with fewer 45-54 year-olds (n=7). Most of us (n=25) spoke a language other than English first (Bengali, Bosnian, Cantonese Chinese, Czech, Finnish, French, German, Hebrew, Italian, Korean, Portuguese, Spanish), although many of us spoke English as a first language (n=18). However, most of us speak English as the usual language now (n=24) or as one of multiple usual languages (n=3), while many of us (n=16) continue to primarily speak other languages (Bengali, Czech, Finnish, French, German, Hebrew, Italian, Portuguese). Most of us originate in either Europe (n=20) or North America (n=15), with n=8 from elsewhere (Australia, Brazil, China, French Polynesia, India, Israel, South Korea). This survey indicates some biases, and in general because we all had access to a specialised conference in our field in person, we recognize that we are a biased subset of many broader communities, and of the movement ecology community in particular.

We understand that not all people are equally represented in movement ecology (or science broadly) and not everyone in movement ecology was part of the in-person conference where this conversation (survey and then discussion) took place. Furthermore, although we invited all conference participants to take the survey, participate in the discussion, and opt-in as an author on this paper, not all in-person meeting participants chose to do so; thus we are a sample even of the conference attendees (see Figure S1 in supplement 2). We grappled with how best to move forward given that we are a subset of the broader movement ecology community. We decided against adding co-authors with identities and perspectives that were missing in the discussion because it would have been a band-aid solution to a more complex issue. Instead, we aimed for transparency in our authorship composition, hoping this manuscript will spur further discussion with a broader range of perspectives.
