## supplement 2 for "Perceived and observed biases within scientific communities: a case study in movement ecology"

### SUPPLEMENT 2: ADDITIONAL RESULTS

**Table S1.** Responses to the open-ended survey question [Q2] 'What do you think the main sources of bias are, if any?', organised into broad categories. Of 135 survey takers, 74 answered this question.

| Bias type | Details | # participants suggesting |
|---|---|---|
| Who does the study | Gender / sexism, race / ethnicity / racism, researcher geography/nationality, class/socioeconomic status, funding disparities by nationality / identity, PI / lab identity / pedigree, cultural, language, institutional, disability, sexual orientation, parental status, age, seniority | 40 |
| What is studied | Taxonomic bias (charisma, size), geography of sites or species, scale and type of movement, terrestrial bias, behaviour type | 26 |
| How it is studied | Conceptual framing, technology reliance/limitations, methods over questions / pattern over process, subjective analysis / study design, small sample sizes and data limitations, theoretical assumptions | 14 |
| How our community works | Access to education / training / data / experience, group think / dogma, academia / structural biases, insular/non-inclusive community, implicit / unconscious bias, networking, ignorance of or ignoring biases, safety / accessibility of work, failed retention, lack of intrafield connections, historical biases, publication / editorial process, trainee treatment, differential impacts of covid, lack of discussions and environmental justice | 25 |

**Figure S1.** Distribution of conference attendees (a) by country of affiliation and (b) by gender.

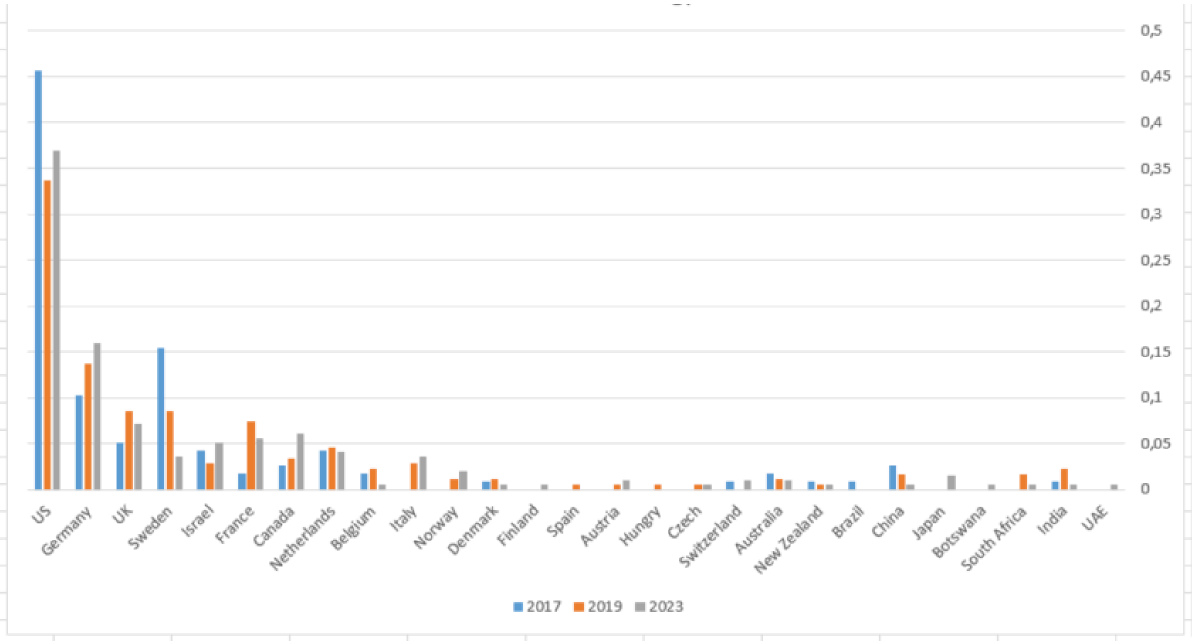


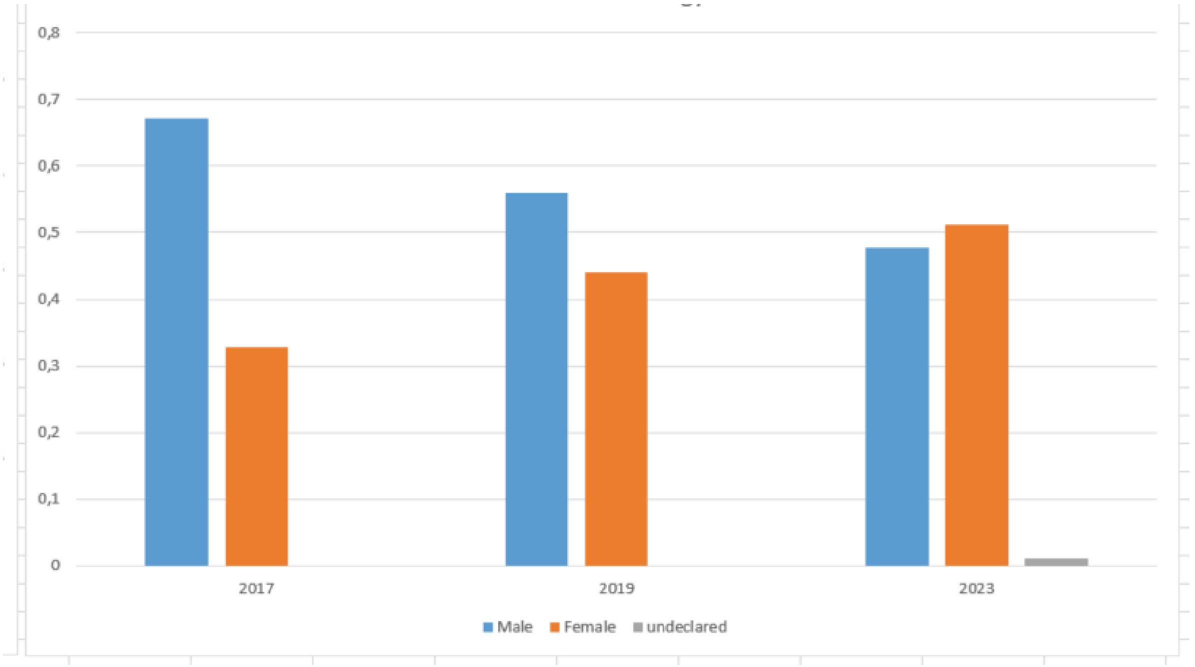

**Figure S2.** The number of times each country was listed as the corresponding author's affiliation for all 370 articles published in the journal *Movement Ecology* (from the first issue in 2013 through January 2023).

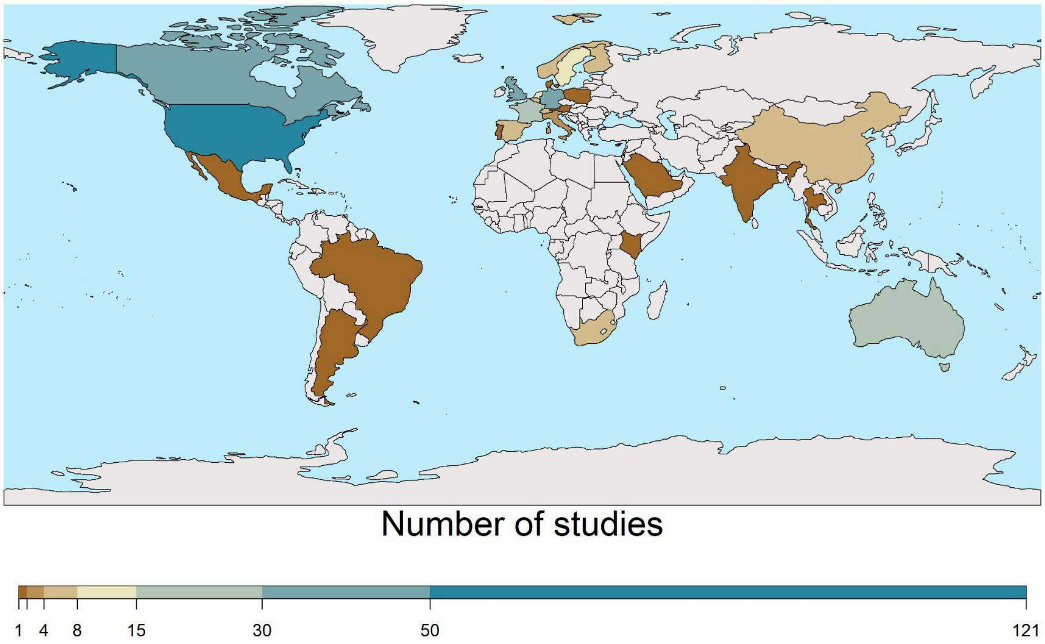

**Figure S3.** Diagnostic plots for the model estimating the relationship between the number of times a country was listed as the first author's first affiliation across all 370 articles published in the journal *Movement Ecology* (from the first issue in 2013 through January 2023) as a function of the country's GDP in that year. The excessively high density of residuals just below zero occurs due to the high number of single publications for a country in one year. The figures were created using the `appraise()` function from the `gratia` package for R (version 0.8.1, Simpson 2023).

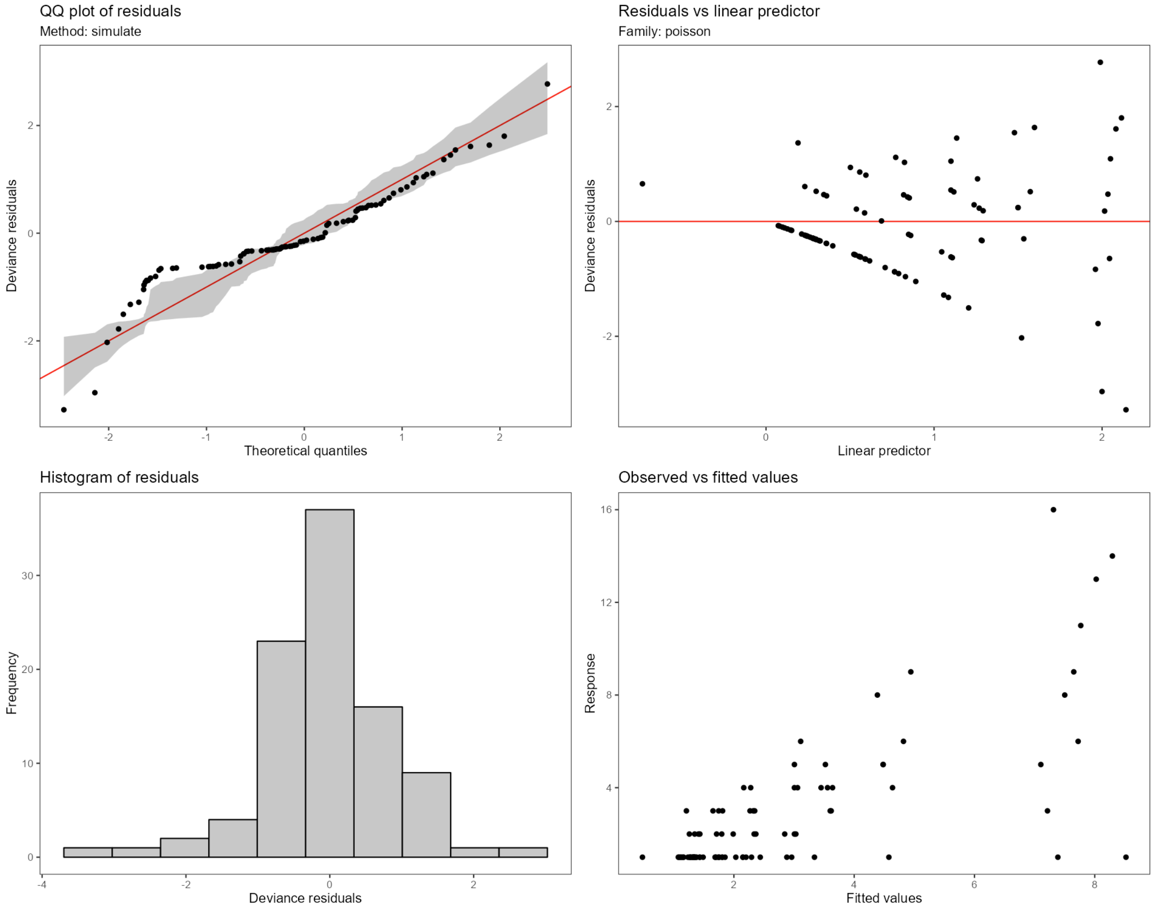

**Figure S4.** Diagnostic plots for the model estimating the relationship between the number of times a country was listed as the first author's first affiliation across the 266 articles with new empirical data published in the journal *Movement Ecology* (from the first issue in 2013 through January 2023) as a function of the country's GDP in that year. The excessively high density of residuals just below zero occurs due to the high number of single publications for a country in one year. The figures were created using the `appraise()` function from the `gratia` package for R (version 0.8.1, Simpson 2023)

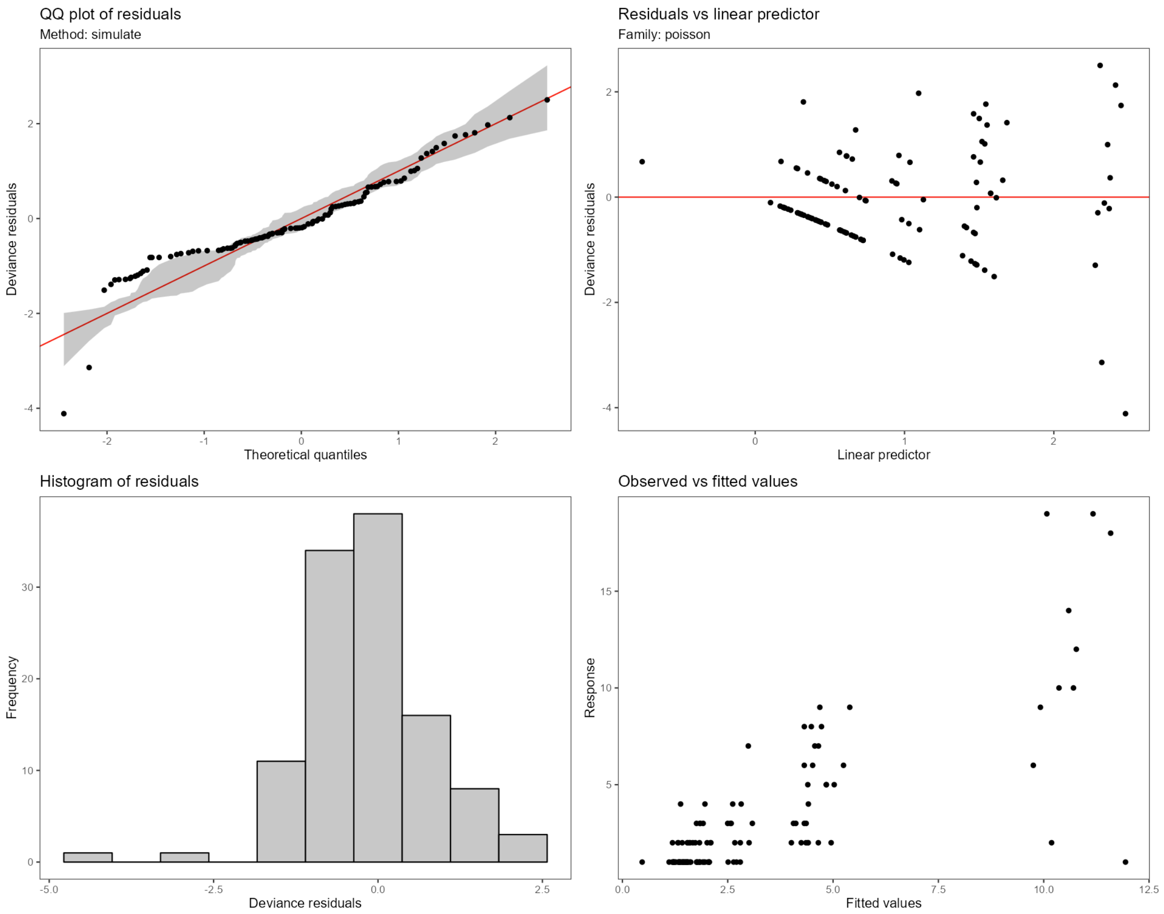
