## supplement 3 for "Perceived and observed biases within scientific communities: a case study in movement ecology"

UNIVERSITY OF MINNESOTA
Driven to Discover®

**Movement Ecology and Bias**

You are invited to be in a research study of perceived biases in the field of Movement Ecology. You were selected as a possible participant because you are registered for the 2023 [redacted] Conference.

This study was developed as an activity within the [redacted] session of the Conference by Francesca Cagnacci (Edmund Mach Foundation, Italy) and Allison Shaw (University of Minnesota, USA). Summarized results of this survey will be presented at the [redacted] during the 2023 [redacted] Conference.

Procedures: If you agree to be in this study, we would ask your opinions on biases within the field of Movement Ecology by completing the survey linked below. The survey will take approximately 15 minutes to complete. When taking the survey, you will be able to take the survey only once and you will not be able to revert back to the previous question or set of questions. Please continue to the end! The responses are meant to be spontaneous with not too much time spent on them.

Confidentiality: Survey responses are anonymous; no identifying data are collected. Any personal information that you share as part of the short response questions will not be shared with others outside of this research study. During the project, information from this study will be kept private and will be stored securely. Information from this research will be used for purposes of this research only and will not be used in future studies or shared with other researchers outside of this specific project.

Voluntary Nature of the Study: Participation in this study is voluntary. Your decision whether or not to participate will not affect your current or future relations with the University of Minnesota, the Edmund Mach Foundation, or the [redacted] If you decide to participate, you are free to not answer any question or withdraw at any time without affecting those relationships.

Contacts and Questions: If you have any questions or concerns about your participation in this study, please contact Allison Shaw or Francesca Cagnacci.

This research has been reviewed and approved by an IRB within the Human Research Protections Program (HRPP). To share feedback privately with the HRPP about your research experience, call the Research Participants' Advocate Line at 612-625-1650 (Toll Free: 1-888-224-8636) or go to z.umn.edu/participants. You are encouraged to contact the HRPP if:
* Your questions, concerns, or complaints are not being answered by the research team.
* You cannot reach the research team.
* You want to talk to someone besides the research team.
* You have questions about your rights as a research participant.
* You want to get information or provide input about this research.

If you agree to participate in this study by taking the survey, please click the arrow below.

---------------------------------------------------- page break ----------------------------------------------------

**Question 1**

To what extent do you see the Movement Ecology community as unbiased? (Where 0 is completely unbiased and 5 is very biased.)

- 0
- 1
- 2
- 3
- 4
- 5

**Question 2**

What do you think the main sources of bias are, if any? Please provide a short answer.

**Question 3**

What is your previous answer *mainly* based on?

- Personal experience
- External evidence (e.g., your readings)
- Speculation

------------------------------------------------- page break -------------------------------------------------

**Question 4**

Which of the following keywords best capture sources of bias in the Movement Ecology community? Please indicate the top three.

| | 1 | 2 | 3 |
|---|---|---|---|
| BIPOC underrepresentation | ○ | ○ | ○ |
| classism | ○ | ○ | ○ |
| dispersal mode bias | ○ | ○ | ○ |
| funding disparities | ○ | ○ | ○ |
| economic policies | ○ | ○ | ○ |
| geographic concentration (of researchers) | ○ | ○ | ○ |
| language | ○ | ○ | ○ |
| methodology | ○ | ○ | ○ |
| perspective | ○ | ○ | ○ |
| racism | ○ | ○ | ○ |
| regional bias (of study sites) | ○ | ○ | ○ |
| sexism | ○ | ○ | ○ |
| study site location | ○ | ○ | ○ |
| taxonomic bias | ○ | ○ | ○ |
| terrestrial environment bias | ○ | ○ | ○ |
| terminology | ○ | ○ | ○ |
| training | ○ | ○ | ○ |

------------------------------------------------- page break -------------------------------------------------

**Question 5**

How do you think the Movement Ecology community could become less biased? Please provide a short answer.

**Question 6**

Which of the following entities should be involved in decreasing bias in the Movement Ecology community? Please indicate the top 3.

| | 1 | 2 | 3 |
|---|---|---|---|
| Individuals | ○ | ○ | ○ |
| Labs | ○ | ○ | ○ |
| Universities | ○ | ○ | ○ |
| Scientific Societies | ○ | ○ | ○ |
| Public funding bodies | ○ | ○ | ○ |
| Private funding bodies | ○ | ○ | ○ |
| Scientific journals | ○ | ○ | ○ |
| Conferences | ○ | ○ | ○ |
| Governments | ○ | ○ | ○ |
| International organizations (e.g., UNEP, IUCN) | ○ | ○ | ○ |
| NGOs | ○ | ○ | ○ |
| Media | ○ | ○ | ○ |
| Social Media | ○ | ○ | ○ |
