## supplement 4 for "Perceived and observed biases within scientific communities: a case study in movement ecology"

**SUPPLEMENT 4: ADDITIONAL METHODS**

To visualize the general patterns of where researchers were affiliated and conducted fieldwork, we created an alluvial plot using data from the 266 studies published in the journa*l Movement Ecology* from its launching in 2013 until January 2023 that collected new empirical data. The plot shows the relative proportion of institutional affiliation by region and links that to the region of fieldwork, while also visualizing the income levels of the fieldwork country. The income levels were determined by country using the 2022 gross national income (GNI) levels from the World Bank Atlas method (The World Bank n.d.). Our labels used for the regional country groupings differed slightly from the World Bank list, both to shorten the amount of text characters in each label and to provide the most succinct and appropriate label for each grouping of countries. The changes made to the region labels include:

1) Australia as a separate region (instead of within East Asia and Pacific) to highlight the amount of studies both based out of Australian institutions and also taking place in Australia.
2) Considering Antarctica a separate region to highlight that the amount of studies conducting fieldwork there (Not a country represented in the World Bank list)
3) Changing 'Middle East and North Africa' to 'Middle East' because all countries represented were from the Middle East and none were from North Africa
4) Changing 'Europe and Central Asia' to 'Europe' because almost all studies were from Western Europe with the exception of Russia, where fieldwork occurred in 3 studies but there were no institutions based there.
5) Changing 'East Asia and Pacific' and 'South Asia' to just 'Asia'. Both of the former region labels had very small sample sizes, so we consolidated them to make it easier to see the relative proportion of studies in this region.
6) Changing 'Sub-Saharan Africa' to 'Africa'. There were no countries in North Africa.
7) Changing 'Latin America & the Caribbean' to ''Latin America'. There were no Caribbean countries

The data for complete information on each study is included in supplement 6.

**References**

The World Bank. (n.d.). The World Bank Atlas method. https://datahelpdesk.worldbank.org/knowledgebase/articles/378832-what-is-the-world-bank-atlas-method..

**Table S2.** Comparison of race/ethnicity categories, showing the ones reported in the datasets on graduate students, postdoctoral researchers and faculty within the United States (all from NCSES) and on the general USA population (from the census), and how we combined the categories for our own analysis. Specifically, NCSES used race and ethnicity as a single axis while the census used race and ethnicity as two separate axes.

<table>
<tr><th>Graduate students and postdocs (NCES)</th><th>Faculty (NCES)</th><th>U.S. population (census)</th><th>Category used for Fig. 5</th></tr>
<tr><td>‘Non-Hispanic, American Indian or Alaska Native’</td><td>Not reported</td><td>‘American Indian or Alaska Native’<br>+ ‘Not Hispanic or Latino’</td><td>‘American Indian and Alaska Native’</td></tr>
<tr><td>‘Non-Hispanic, Asian’</td><td>‘Non-Hispanic, Asian’</td><td rowspan="2">‘Asian or Pacific Islander’<br>+ ‘Not Hispanic or Latino’</td><td rowspan="2">‘Asian (including Pacific Islander)’</td></tr>
<tr><td>‘Non-Hispanic, Native Hawaiian or Other Pacific Islander’</td><td>Not reported</td></tr>
<tr><td>‘Non-Hispanic, black or African American’</td><td>‘Non-Hispanic, black’</td><td>‘Black or African American’<br>+ ‘Not Hispanic or Latino’</td><td>‘Black and African American’</td></tr>
<tr><td rowspan="4">‘Hispanic or Latino’</td><td rowspan="4">‘Hispanic, any race’</td><td>‘American Indian or Alaska Native’<br>+ ‘Hispanic or Latino’</td><td rowspan="4">‘Hispanic and Latino’</td></tr>
<tr><td>‘Asian or Pacific Islander’<br>+ ‘Hispanic or Latino’</td></tr>
<tr><td>‘Black or African American’<br>+ ‘Hispanic or Latino’</td></tr>
<tr><td>‘White’<br>+ ‘Hispanic or Latino’</td></tr>
<tr><td>‘Non-Hispanic, white’</td><td>‘Non-Hispanic, white’</td><td>‘White’</td><td>‘White’</td></tr>
</table>

| | | | |
|---|---|---|---|
| | | + 'Not Hispanic or Latino' | |
| 'Non-Hispanic, more than one race' | 'Non-hispanic, other races including multiracial individuals' | *Not reported* | *Not plotted* |
