## supplement 5 for "Perceived and observed biases within scientific communities: a case study in movement ecology"

### [redacted] Discussion topic: Is the Movement Ecology community unbiased?

#### BACKGROUND: WHAT IS THIS ABOUT?

•**The** [redacted] is designed to address ways to improve diversity and inclusion in science by providing a safe environment for informal and meaningful conversations amongst colleagues of all career stages.
The program supports the professional growth of all members of our communities.
•Previous [redacted] dealt with general sources of inequality, e.g. sex-bias in science.
•**Movement Ecology** discipline and community is now mature enough to hold responsibility for asking these questions **within its domain**.
•We formulated some simple questions for a survey, which was submitted to [redacted] participants before the meeting.
•In parallel, we assessed some **simple objective metrics of bias**, taking as a reference the papers published in the **Movement Ecology journal**.
•The aim of this handout is to summarise the results of the survey and bias assessment, to be used as a basis for discussion at the [redacted]
•**Please participate in the** [redacted] **meeting!! We would value hearing your ideas to help shape next steps.**

#### SURVEY RESULTS

**1. To what extent do you see the Movement Ecology community as unbiased?**
(0 = completely unbiased; 5 = very biased.)

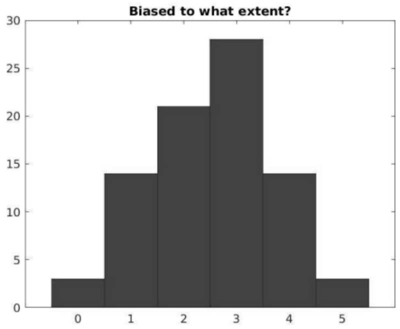


**2. What is your previous answer mainly based on?**

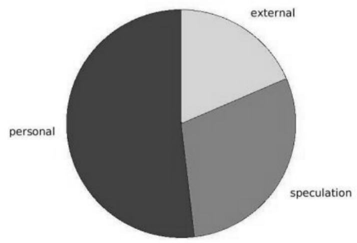


**3. Which of the following keywords best capture sources of bias in the Movement Ecology community?**
(The number of times each response was chosen in the top 3).

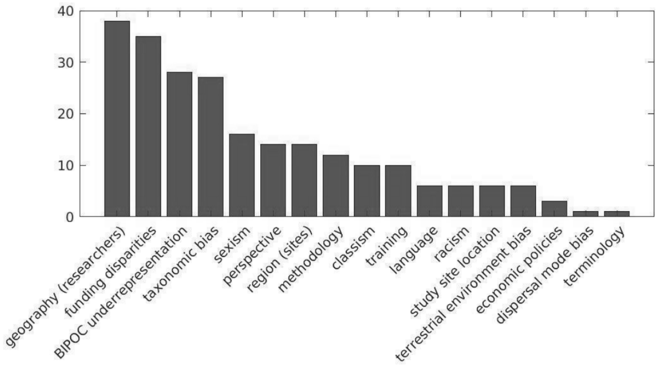


**4. Which of the following entities should be involved in decreasing bias in the Movement Ecology community?**
(The number of times each response was chosen in the top 3).

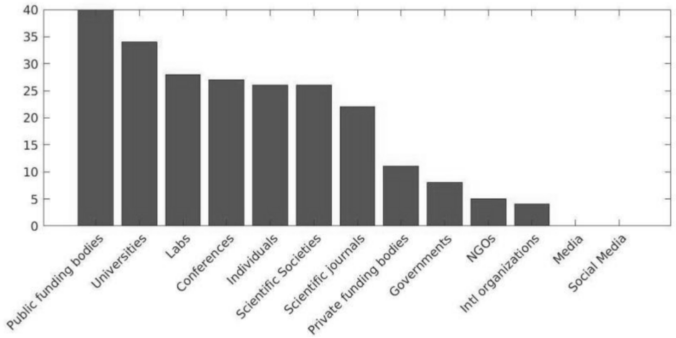

**5. How do you think the Movement Ecology community could become less biased?**

| Strategy | Propositions | N° Survey takers | Tot n° |
|---|---|---|---|
| Financial | • Funding<br>• Geography-specific<br>• Stage-specific (e.g. ECR)<br>• Group-specific ((e.g. historically marginalized<br>• Cost reduction (research, equipment, pubblication) | 20<br>8<br>4<br>3<br>3 | 38 |
| Personnel | • Training opportunities (target by country, target underrappresented groups)<br>• Collaboration/Knowledge sharing (across fields, taxa, perspectives, regions, communities, indigenous perspectives)<br>• Elevate researchers from under-represented backgrounds<br>• Better advising (more inclusive, aknowledge trainees) | 12 (5, 4)<br>6<br>4<br>2 | 24 |
| Scientific process | • Methodological approaches (stronger framing + less descriptive methods, technological advances, make data avilable, more observation, synthesis, targeted experiments)<br>• Changing norms (gatekeeping, pubblication prestige, what is founded, pubblication lenguage)<br>• Systems studied (increase geographic and taxa diversity)<br>• Pubblication process ( reduce bias in review, more geographic opportunities) | 9 (3, 2)<br>5<br>5<br>5 | 24 |
| Events | • Discussion/learning opportunities (unconscious bias, how to shift culture)<br>• Conferences (alternatives to [redacted] or reduce disparity at [redacted])<br>• Learning (study our biases, learn history of our science)<br>• Networking<br>• Special issues in movement ecology<br>• Workshops (technical) | 4<br>2<br>2<br>1<br>1<br>1 | 11 |

#### RESULTS FROM ANALYZING MOVEMENT ECOLOGY ARTICLES

Authorship data from 370 papers in Movement Ecology from the first issue through Jan 2023, looking at the first country listed as an affiliation of each paper's first author. Then plotting the number of first author papers by country versus that country's GDP.

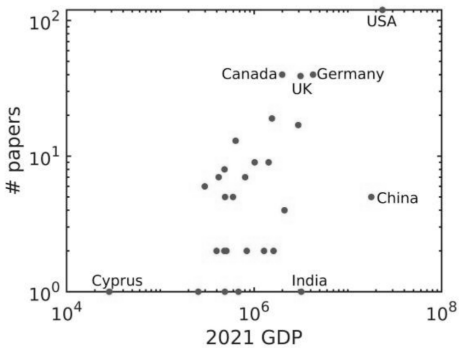


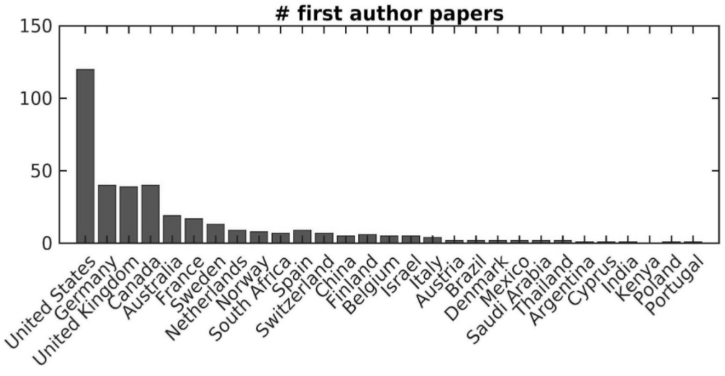


#### OTHER SOURCES OF BIAS: RESULTS FROM U.S. DEMOGRAPHY

The proportion of the total population (y-axis) for faculty (fac), postdocs (pd), graduate students (grad; US citizens and permanent residents only), and the general U.S. population (USA)

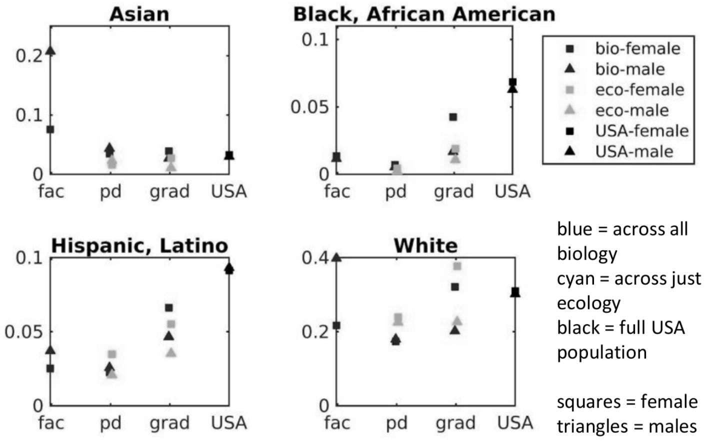
